# dbGuide: A database of functionally validated guide RNAs for genome editing in human and mouse cells

**DOI:** 10.1101/2020.08.24.253245

**Authors:** Alexander A. Gooden, Christine N. Evans, Timothy P. Sheets, Michelle E. Clapp, Raj Chari

## Abstract

With the technology’s accessibility and ease of use, CRISPR has been employed widely in many different organisms and experimental settings. As a result, thousands of publications have used CRISPR to make specific genetic perturbations, establishing in itself a resource of validated guide RNA sequences. While numerous computational tools to assist in the design and identification of candidate guide RNAs exist, these are still just at best predictions and generally, researchers inevitably will test multiple sequences for functional activity. Here, we present *dbGuide* (https://sgrnascorer.cancer.gov/dbguide), a database of functionally validated guide RNA sequences for CRISPR/Cas9-based knockout in human and mouse. Our database not only contains computationally determined candidate guide RNA sequences, but of even greater value, over 4000 sequences which have been functionally validated either through direct amplicon sequencing or manual curation of literature from over 1000 publications. Finally, our established framework will allow for continual addition of newly published and experimentally validated guide RNA sequences for CRISPR/Cas9-based knockout as well as incorporation of sequences from different gene editing systems, additional species, and other types of site-specific functionalities such as base editing, gene activation, repression, and epigenetic modification.

## INTRODUCTION

CRISPR/Cas9-based genome editing has been an indispensable technology for understanding the biology of living organisms^1,2^. As a result, a tremendous effort has been invested in aiding researchers in designing and executing CRISPR-based experiments by providing critical resources such as reagents and protocols, and software tools that allow a user to rapidly identify guide RNA sequences with specific predicted on-target and off-target characteristics^3–18^ (Table 1). However, these sequences are essentially predictions, often requiring assessment of multiple candidate guide RNAs.

**Table 1:**
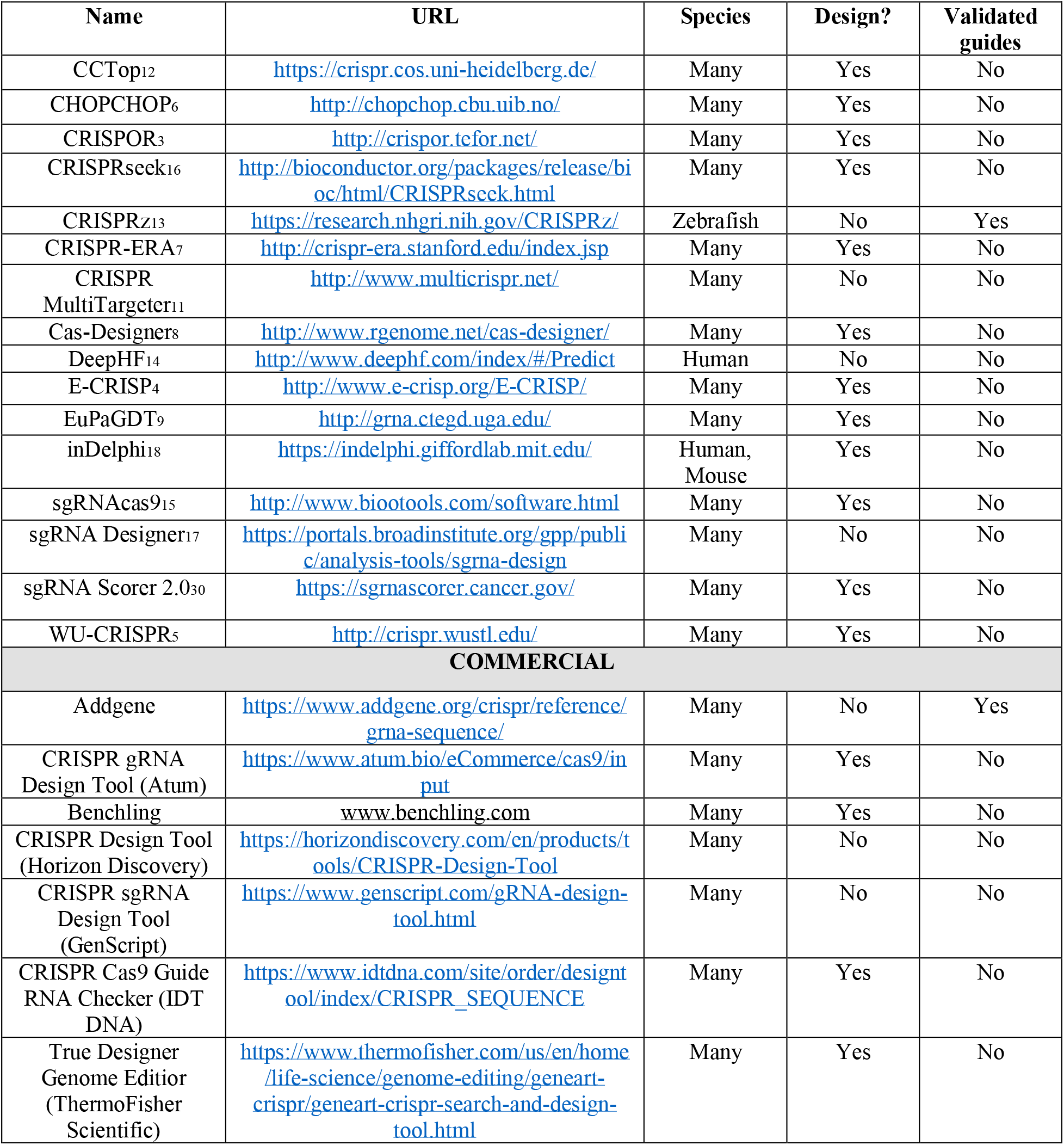
List of sgRNA design tools

Over the last seven years, there has been an explosion of literature utilizing CRISPR/Cas9 to knockout specific genes or genomic regions, of which the majority is in human or mouse model systems. The culmination of all of this work has provided an opportunity to mine this information for guide RNA sequences shown to be functional at either a genotypic and/or phenotypic level. Variation in reporting standards, typographical errors, and relegation of sequences into supplementary materials are amongst a few reasons why curation of this information has been non-trivial.

To this end, we present *dbGuide*, a database of functionally validated guide RNA sequences for CRISPR/Cas9 mediated knockout experiments using human or mouse cells (http://sgrnascorer.cancer.gov/dbguide). We have manually curated guide RNA sequences from over 1000 peer-reviewed articles with each sequence having direct reference to the original publication. We are also making results of targeted amplicon sequencing for ~2000 unique sgRNA sequences tested individually in human (293T) or mouse (NIH-3T3 or P19) cultured cells publicly searchable in our database. In total, these efforts encompass nearly 6000 unique guide RNA sequences for which some level of validated activity exists. To our knowledge, this represents the largest database of functionally validated guide RNA sequences. We expect this to be a continually growing resource through multiple mechanisms. We have provided a downloadable template to encourage researchers to submit their newly published/validated sequences and our computational framework will also allow for the inclusion of sequences for CRISPR systems other than for *S. pyogenes*, modalities such as base editing, gene activation/repression and epigenetic modifications, and sequences used in other species.

## DATA COLLECTION

### Master list of human and mouse sgRNA sequences

In addition to functionally validated sgRNA sequences, in order to provide further utility to the database, computationally designed sequences were also obtained from a variety of sources, primarily focused on protein-coding genes^17,19–26^. These sources are listed in Supplementary Table 1.

### Published sgRNA sequences

A broad search of the PubMed database for “CRISPR OR Cas9” was performed and yielded over 15,000 indexed citations. Subsequently, review articles, publications not using human or mouse cells, publications not using *S. pyogenes* Cas9, publications not performing knock-out experiments were excluded. In total, guide RNA sequences were sourced from a total of 1322 peer-reviewed articles (Supplementary Table 2).

### Targeted amplicon sequencing data

Quantitative editing data for nearly 2000 sgRNA sequences were generated internally from either transfection of Cas9/sgRNA plasmids (1 ug) or Cas9 protein (4 ug) / *in vitro* transcribed (IVT) (2.25 ug) RNA into mouse (NIH-3T3 or P19) or human (HEK293T) cells in a 24-well format. Cas9 protein was produced using plasmid Addgene-62731, a gift from Niels Geijsen (Addgene plasmid # 62731; http://n2t.net/addgene:62731; RRID:Addgene_62731)^27^. IVT RNA was produced using a similar protocol as previously published^28^. Cas9/RNP complexing and transfection using Lipofectamine 2000 were also performed similarly to as previously published^29^. PCR from genomic DNA was performed and amplicons were sequenced using the Illumina MiSeq V2 300 cycle kit using the PE (2 × 150) format. List of all sgRNAs sequences tested are listed in Supplementary Table 3.

## DATA PROCESSING

### Mapping information and on/off-target scoring metrics

For all sgRNA sequences obtained, genomic locations of the corresponding target sites were obtained/verified using UCSC BLAT against either the hg38 or mm10 reference genomes. Subsequently, sgRNA locations were cross-referenced with the Gencode V32 (human) or Gencode VM23 (mouse) gene/transcript annotations to determine which transcript(s) each sgRNA could target.

For on-target metrics, *sgRNA Scorer 2.0*^30^, Rule Set 2^17^, and *FORECasT*^31^ scores were downloaded/calculated for each sgRNA and if a score could not be obtained, a value of “NV” was denoted. For off-target analysis, *Guidescan* 1.0^32^ values were generated for each guide and similarly, for those guides for which a score could not be obtained, a value of “NV” was given.

### Analysis of targeted amplicon sequencing data

Paired end raw FASTQ files were merged using *FLASH*^33^, filtered for low quality bases, subsequently mapped to the designated genomic locations in hg38/mm10 using bwa mem, and then sorted and indexed bam files were generated using samtools^34^. A custom python snakemake^35^ pipeline was made to calculate non-homologous end joining (NHEJ) mutation frequencies. This analysis pipeline is publicly available at https://github.com/rajchari2/ngs_amplicon_analysis.

## IMPLEMENTATION AND ACCESS

*dbGuide* uses a simple HTML interface which utilizes the *datatables* and *highcharts* javascript libraries for displaying data in tabular and graphical formats, respectively (Figure 1A). The application is built in python using django with a MySQL database used for data storage and retrieval. The database can be accessed without e-mail registration or login.

**Figure 1.**
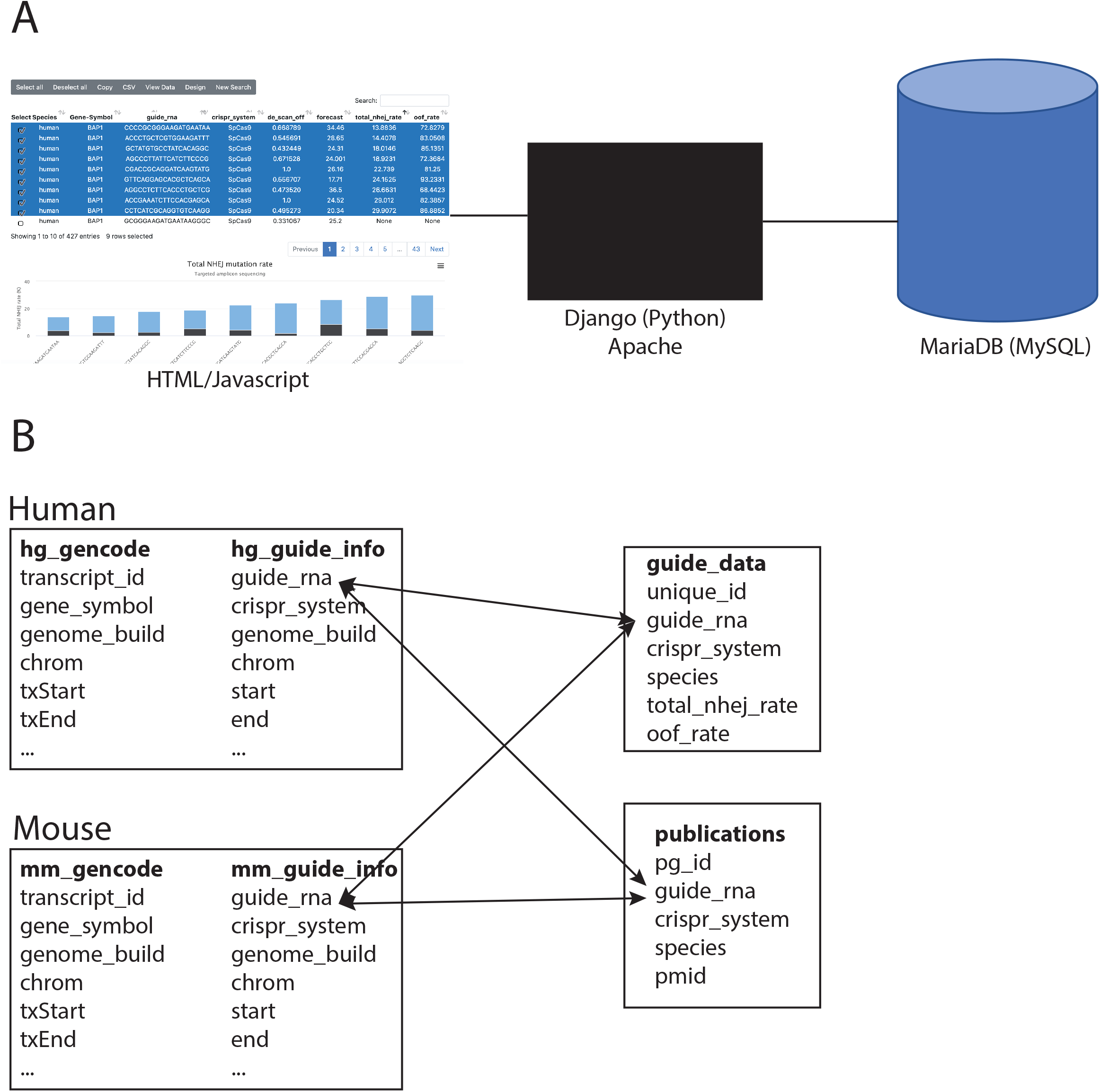
Structure of the *dbGuide* database. **(A)** Components of the *dbGuide* database. The user interface was developed using html and javascript and the application is managed on an Apache webserver using django. All underlying data is stored in a MariaDB (MySQL) database. **(B)** Schema of the MySQL database. Each species, currently limited to human and mouse, has a table of sgRNA sequences with pre-computed metrics and a table with gene annotation information. sgRNA sequences obtained from publications are stored in a single table and the “species” field is used to determine which species the guide RNA was used. Similarly, sequences from targeted amplicon sequencing data also have the “species” field for this purpose.

Within the MySQL relational database, for both human and mouse, there was central sgRNA table with genome target position and all metrics pre-calculated and a gene annotation table which has the location of all protein coding genes based on *Gencode* annotation. Finally, there are two separate tables for summarizing the amplicon sequencing data and the publication-validated sequences. A depiction of the schema is shown in Figure 1B.

## DATABASE CONTENTS AND FEATURES

### Opening user interface

The introductory user interface is very simple (Figure 2A). The user first specifies whether to search in the human or mouse genome and then can provide either a chromosomal position (in BED format), gene symbol, or *Ensembl* gene or transcript ID. In addition, a link to an excel spreadsheet template is provided for researchers wishing to contribute to the database.

**Figure 2.**
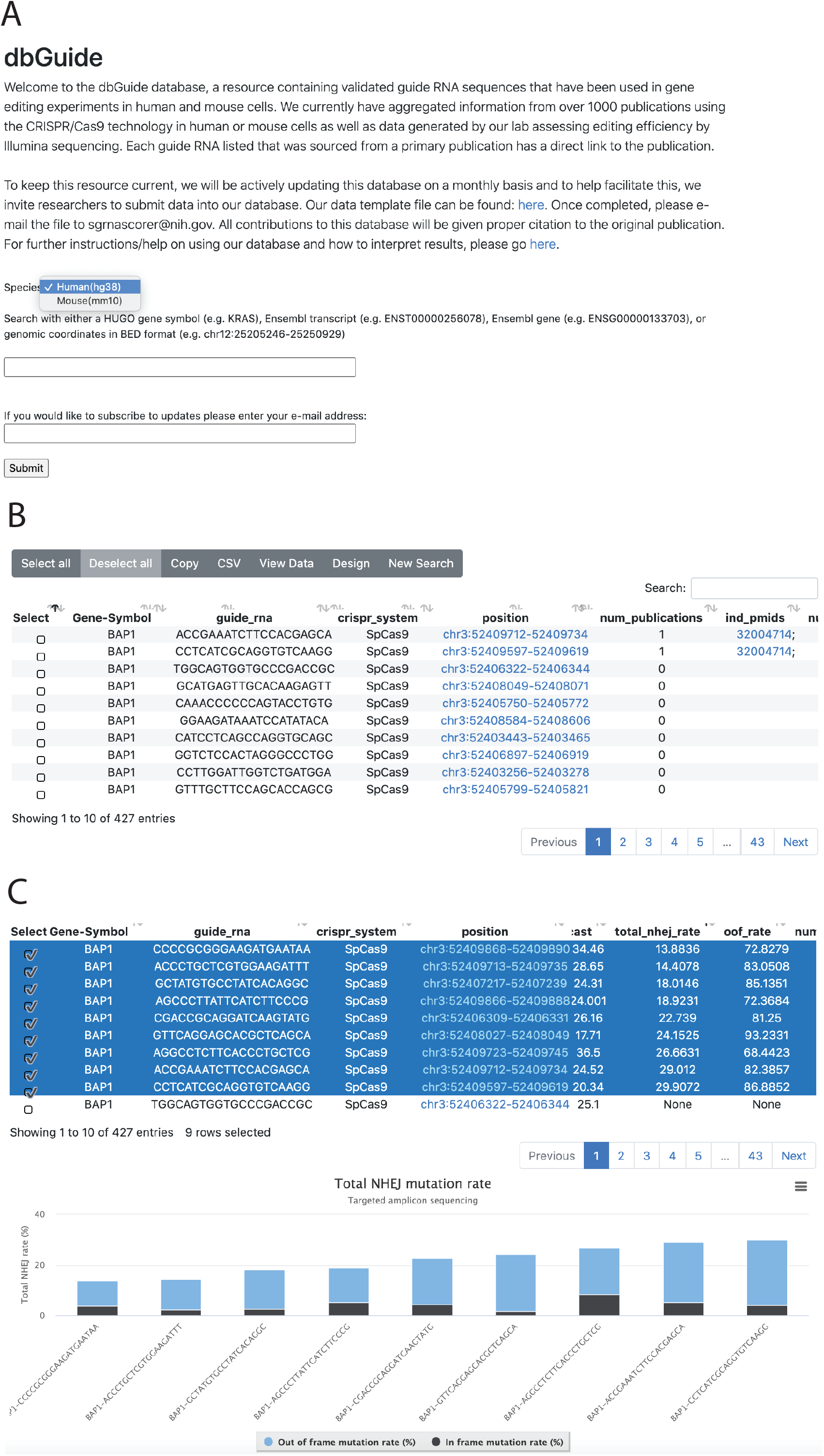
Screen shots of the key user interfaces. **(A)** Opening window of the *dbGuide* database. The user must first select whether to search within the human/mouse genome and then can specify a HGNC gene symbol, chromosomal coordinate in BED format, or ENSEMBL gene/transcript ID. **(B)** Window depicting results of a search by gene symbol. Selectable rows of sgRNA sequences are returned with various pre-computed metrics and PubMed identifiers, if the sequence had been used in a peer-reviewed publication. **(C)** Graphical representation of sgRNA sequences for which targeted amplicon sequencing data were generated. Stacked bar plots display the percentage of non-homologous end joining (NHEJ) mutation events that results in an “in-frame” (black) or “out of frame” (blue) amino acid change.

### Search results when searching by gene/coordinate/accession

Upon entering a valid search term, the user will be shown a table of results detailing all of the guide RNA sequences identified based on the search criteria (Figure 2B). The table displays the following columns:

**Gene-Symbol** – most recent official gene symbol for a gene,
**guide_rna** – sgRNA spacer sequence (without PAM),
**crispr_system** - Cas9 ortholog, currently restricted to SpCas9,
**position** - genomic location of the sgRNA target site in either human (hg38) or mouse (mm10) assembly
**in_protein_coding_exon** - whether the guide RNA targets a protein coding exon,
**num_transcripts** - the number of GENCODE transcripts targeted,
**transcript_id_list** - list of GENCODE transcripts (by ENSEMBL ID),
**sgrnascorer** - Predicted activity using the sgRNA Scorer 2.0 algorithm (−3 to 3). Higher the value, greater the predicted activity,
**rule_set_2** - Predicted activity using the Rule Set 2 algorithm (0 to 1). Higher the value, greater the predicted activity.
**guide_scan_off** – Guidescan off-target score (0 to 1). Higher the value, higher the specificity.
**forecast** – Favored Outcomes of Repair Events at Cas9 targets (FORECasT) score for predicting mutational outcomes (in frame indel %, lower values preferred for knockouts),
**total_nhej_rate** – if amplicon sequencing data exists, value between 0 and 100, or “None” otherwise
**oof_rate** – out of frame mutation rate which is the proportion of mutated reads which would lead to an out of frame mutation
**num_publications** – number of published articles using the selected guide RNA sequence
**ind_pmids –** list of articles by PubMed ID which used selected guide RNA sequence
**num_screens** – number of pooled CRISPR screens in which the guide RNA sequences was enriched or depleted
**screen_pmids** – list of articles by PubMed ID of CRISPR-based pooled screens in which this guide was enriched or depleted
**sources** – list of sources from which the guide RNA sequence has been curated from

### Viewing sequence editing data

In addition to sorting by total NHEJ rate, one can view the data in a graphical format. By selecting sgRNA sequences for which data exists, the user can click the “View Data” which will show a stacked bar plot breaking down the total NHEJ rate between “in frame” and “out of frame” percentage (Figure 2C). This can be specifically important for knockout experiments where a higher out of frame (OOF) mutation percentage gives higher probability of protein loss.

### Downloading sequences for use in experiments

Upon identifying guide RNA sequences of interest, the user can export the sequences in a “ready to order” format by clicking on the “Design”. A tab-delimited text file is generated listing oligonucleotides needed for either ligation into a plasmid or *in vitro* transcription as well as a PDF protocol describing what compatible vectors can be used and how they can be obtained.

## DATA AVAILABILITY

Sorted BAM files for amplicon sequencing data are available upon reasonable request.

## SUPPLEMENTARY DATA

Supplementary Table 1 – List of sources for sgRNA sequences

Supplementary Table 2 – List of sgRNA sequences collected from published literature

Supplementary Table 3 – List of sgRNA sequences with amplicon sequencing data

## ACKNOWLEDGEMENTS

We would like to acknowledge Troy Taylor, William Gillette, Jane Jones, and Dominic Esposito for kindly providing Cas9 protein. We would also like to acknowledge the Biomedical Informatics and Data Science (BIDS) and Frederick Research Computing Environment (FRCE) for providing assistance in web hosting and computational support.

## REFERENCES

1. Mali, P. et al. RNA-guided human genome engineering via Cas9. Science 339, 823–826 (2013).

2. Cong, L. et al. Multiplex genome engineering using CRISPR/Cas systems. Science 339, 819–823 (2013).

3. Haeussler, M. et al. Evaluation of off-target and on-target scoring algorithms and integration into the guide RNA selection tool CRISPOR. Genome Biol. 17, 148 (2016).

4. Heigwer, F., Kerr, G. & Boutros, M. E-CRISP: fast CRISPR target site identification. Nat. Methods 11, 122–123 (2014).

5. Hiranniramol, K., Chen, Y., Liu, W. & Wang, X. Generalizable sgRNA design for improved CRISPR/Cas9 editing efficiency. Bioinforma. Oxf. Engl. 36, 2684–2689 (2020).

6. Labun, K. et al. CHOPCHOP v3: expanding the CRISPR web toolbox beyond genome editing. Nucleic Acids Res. 47, W171–W174 (2019).

7. Liu, H. et al. CRISPR-ERA: a comprehensive design tool for CRISPR-mediated gene editing, repression and activation. Bioinforma. Oxf. Engl. 31, 3676–3678 (2015).

8. Park, J., Bae, S. & Kim, J.-S. Cas-Designer: a web-based tool for choice of CRISPR-Cas9 target sites. Bioinforma. Oxf. Engl. 31, 4014–4016 (2015).

9. Peng, D. & Tarleton, R. EuPaGDT: a web tool tailored to design CRISPR guide RNAs for eukaryotic pathogens. Microb. Genomics 1, e000033 (2015).

10. Pliatsika, V. & Rigoutsos, I. ‘Off-Spotter’: very fast and exhaustive enumeration of genomic lookalikes for designing CRISPR/Cas guide RNAs. Biol. Direct 10, 4 (2015).

11. Prykhozhij, S. V., Rajan, V., Gaston, D. & Berman, J. N. CRISPR multitargeter: a web tool to find common and unique CRISPR single guide RNA targets in a set of similar sequences. PloS One 10, e0119372 (2015).

12. Stemmer, M., Thumberger, T., Del Sol Keyer, M., Wittbrodt, J. & Mateo, J. L. CCTop: An Intuitive, Flexible and Reliable CRISPR/Cas9 Target Prediction Tool. PloS One 10, e0124633 (2015).

13. Varshney, G. K. et al. CRISPRz: a database of zebrafish validated sgRNAs. Nucleic Acids Res. 44, D822–826 (2016).

14. Wang, D. et al. Optimized CRISPR guide RNA design for two high-fidelity Cas9 variants by deep learning. Nat. Commun. 10, 4284 (2019).

15. Xie, S., Shen, B., Zhang, C., Huang, X. & Zhang, Y. L. sgRNAcas9: a software package for designing CRISPR sgRNA and evaluating potential off-target cleavage sites. PloS One 9, e100448 (2014).

16. Zhu, L. J., Holmes, B. R., Aronin, N. & Brodsky, M. H. CRISPRseek: a bioconductor package to identify target-specific guide RNAs for CRISPR-Cas9 genome-editing systems. PloS One 9, e108424 (2014).

17. Doench, J. G. et al. Optimized sgRNA design to maximize activity and minimize off-target effects of CRISPR-Cas9. Nat. Biotechnol. 34, 184–191 (2016).

18. Shen, M. W. et al. Predictable and precise template-free CRISPR editing of pathogenic variants. Nature 563, 646–651 (2018).

19. Doench, J. G. et al. Rational design of highly active sgRNAs for CRISPR-Cas9-mediated gene inactivation. Nat. Biotechnol. 32, 1262–1267 (2014).

20. Sanjana, N. E., Shalem, O. & Zhang, F. Improved vectors and genome-wide libraries for CRISPR screening. Nat. Methods 11, 783–784 (2014).

21. Hart, T. et al. High-Resolution CRISPR Screens Reveal Fitness Genes and Genotype-Specific Cancer Liabilities. Cell 163, 1515–1526 (2015).

22. Chari, R., Mali, P., Moosburner, M. & Church, G. M. Unraveling CRISPR-Cas9 genome engineering parameters via a library-on-library approach. Nat. Methods 12, 823–826 (2015).

23. Wang, T. et al. Gene Essentiality Profiling Reveals Gene Networks and Synthetic Lethal Interactions with Oncogenic Ras. Cell 168, 890–903.e15 (2017).

24. Wang, T. et al. Identification and characterization of essential genes in the human genome. Science 350, 1096–1101 (2015).

25. Tzelepis, K. et al. A CRISPR Dropout Screen Identifies Genetic Vulnerabilities and Therapeutic Targets in Acute Myeloid Leukemia. Cell Rep. 17, 1193–1205 (2016).

26. Henriksson, J. et al. Genome-wide CRISPR Screens in T Helper Cells Reveal Pervasive Crosstalk between Activation and Differentiation. Cell 176, 882–896.e18 (2019).

27. D’Astolfo, D. S. et al. Efficient intracellular delivery of native proteins. Cell 161, 674–690 (2015).

28. Richardson, C. D., Ray, G. J., Bray, N. L. & Corn, J. E. Non-homologous DNA increases gene disruption efficiency by altering DNA repair outcomes. Nat. Commun. 7, 12463 (2016).

29. Kim, K. et al. Genome surgery using Cas9 ribonucleoproteins for the treatment of age-related macular degeneration. Genome Res. 27, 419–426 (2017).

30. Chari, R., Yeo, N. C., Chavez, A. & Church, G. M. sgRNA Scorer 2.0: A Species-Independent Model To Predict CRISPR/Cas9 Activity. ACS Synth. Biol. 6, 902–904 (2017).

31. Allen, F. et al. Predicting the mutations generated by repair of Cas9-induced double-strand breaks. Nat. Biotechnol. (2018) doi:10.1038/nbt.4317.

32. Perez, A. R. et al. GuideScan software for improved single and paired CRISPR guide RNA design. Nat. Biotechnol. 35, 347–349 (2017).

33. Magoč, T. & Salzberg, S. L. FLASH: fast length adjustment of short reads to improve genome assemblies. Bioinforma. Oxf. Engl. 27, 2957–2963 (2011).

34. Li, H. et al. The Sequence Alignment/Map format and SAMtools. Bioinforma. Oxf. Engl. 25, 2078–2079 (2009).

35. Köster, J. & Rahmann, S. Snakemake-a scalable bioinformatics workflow engine. Bioinforma. Oxf. Engl. 34, 3600 (2018).

